# Population genomics of *C. melanopterus* using target gene capture data: demographic inferences and conservation perspectives

**DOI:** 10.1101/068106

**Authors:** Pierpaolo Maisano Delser, Shannon Corrigan, Matthew Hale, Chenhong Li, Michel Veuille, Serge Planes, Gavin Naylor, Stefano Mona

## Abstract

Population genetics studies on non-model organisms typically involve sampling few markers from multiple individuals. Next-generation sequencing approaches open up the possibility of sampling many more markers from fewer individuals to address the same questions. Here, we applied a target gene capture method to deep sequence ∼1000 independent autosomal regions of a non-model organism, the blacktip reef shark (*Carcharhinus melanopterus*). We devised a sampling scheme based on the predictions of theoretical studies of metapopulations to show that sampling few individuals, but many loci, can be extremely informative to reconstruct the evolutionary history of species. We collected data from a single deme (SID) from Northern Australia and from a scattered sampling representing various locations throughout the Indian Ocean (SCD). We explored the genealogical signature of population dynamics detected from both sampling schemes using an ABC algorithm. We then contrasted these results with those obtained by fitting the data to a non-equilibrium finite island model. Both approaches supported an *Nm* value ∼40, consistent with philopatry in this species. Finally, we demonstrate through simulation that metapopulations exhibit greater resilience to recent changes in effective size compared to unstructured populations. We propose an empirical approach to detect recent bottlenecks based on our sampling scheme.

## Introduction

Obtaining sufficient sequence information is no longer a limiting factor for deciphering the demographic and selective history of a species. Many studies now use population genomic approaches to shed light on evolutionary questions that also have practical applications for species conservation and management. While next-generation sequencing (NGS) techniques have made such approaches possible, studying non-model organisms without a reference genome remains a challenge. Thus far, whole-genome re-sequencing^1^, transcriptome analysis^2^ and *de novo* restriction site associated DNA (RAD) sequencing^3^ have been used to investigate the population dynamics of non-model organisms. All of these approaches have limitations: re-sequencing whole genomes of many individuals remains expensive and superfluous when reliable results can be obtained using a subset of the genome^4^. Generating transcriptome data sets suitable for investigating adaptive change can be complicated by the requirement for high quality RNA and sampling equivalent life stages and tissue types for all individuals^5^. Short sequence lengths, high rates of allelic dropout^6^ and missing data^7^, as well as the vagaries of the bioinformatics pipeline that is used^8^, can influence estimates of genetic diversity obtained with RAD-seq approaches. Target gene capture approaches offer a promising alternative to these methods. They permit sequencing of pre-selected regions of the genome (see the review from^9^) that have been determined *a priori* to be informative for the research question and are generally effective across a range of sample types and qualities. This allows the generation of relatively unbiased, more complete datasets than can be obtained using RAD technologies, and at an affordable cost compared with whole-genome re-sequencing.

The population structure of the study organism is another factor that may complicate our ability to make robust demographic inferences from population genomic data. Previous work has shown that ignoring metapopulation structure can sometimes mislead interpretations of the demographic and evolutionary history of a species^10-13^. This is a problem because populations are rarely entirely isolated in nature but instead belong to a network of demes that exchange migrants at different rates. Despite these concerns, few studies have attempted to estimate demographic parameters under complex metapopulation models (see^14^ and ^15^ for some exceptions). The predicted coalescent patterns that are associated with metapopulations sampled at different levels are of special interest. The gene genealogy of a metapopulation is characterized by the product of the effective population size N and the migration rate *m*^16-19^. The coalescent history of a sample of lineages can be divided into two phases: the *scattering* phase within each single deme and the *collecting* phase when each of the lineages comes from a different deme^16,20^. This separation of time scales holds true for several metapopulation models, including range expansions^19^. Contrasting the shape of the genealogy at different sampling levels (e.g. “single deme” vs. “scatter”, where each lineage comes from a different deme) indirectly provides information on the long term *Nm*^18^. It is therefore possible to make inferences about the evolutionary history of the metapopulation using relatively simple demographic models that investigate changes in effective population size across specific sampling schemes. A few individuals judiciously sampled from the range distribution of a species can be sufficient to uncover its demography, when enough loci are available for statistical inferences. Such an approach is now tenable with the large datasets that are being generated using NGS and has important applications for the study of endangered animals, or of any organism for which specimens are not easily obtained.

Here, we demonstrate the use of such an approach to study the demographic history of a non-model organism, the blacktip reef shark (*Carcharhinus melanopterus*). These animals are highly philopatric and are often found on remote coral islands and atolls^21,22^. Their proclivity for philopatry and strong habitat preference for coral reefs predisposes them to exhibit a disjunct meta-population structure over their widespread distribution throughout the Indo-Pacific. We devised a two-layered sampling scheme based on the above theoretical considerations associated with metapopulations^20^: we collected individuals from a single deme from Northern Australia and from a scattered sampling from various locations throughout the Indo-Pacific. We then used a newly developed target gene capture approach^23^ to generate sequence data for ~1000 pre-specified independent orthologous autosomal regions. This approach allowed us to minimise the sampling of individuals, while ensuring a comprehensive sampling of loci for a species for which no closely related reference genome exists. Demographic parameters were estimated in two ways: i) indirectly by contrasting the gene genealogies at different sampling levels in the metapopulation (“single deme” vs. “scatter”); ii) directly by applying a non-equilibrium finite island model to both sampling schemes. We developed an Approximate Bayesian Computation (ABC) framework^24,25^ with recombination to estimate parameters and to compare demographic models. The site frequency spectrum (SFS) computed on unphased data was used as summary statistic, thus avoiding the level of uncertainty that is introduced by phasing and haplotype reconstruction.

*Carcharhinus melanopterus* inhabits reef flats and sheltered lagoons^21,22^in the Indian and Pacific Oceans (the “Indo-Pacific”). While their biology has been widely studied their population dynamics and dispersal patterns are largely unknown. Nevertheless, insights into their movements and population health status can be gained through assessments of their evolutionary history. The species is considered globally near threatened (NT) by the International Union for Conservation of Nature (IUCN) Red list, with a decreasing population^26^. *Carcharhinus melanopterus* appears to be locally abundant in many areas but recent fishing and anthropogenic pressures on reef environments may have affected their distribution and population dynamics^26-28^. A recent bottleneck was detected in Moorea^29^ suggesting that decreases in effective population size may also have occurred in some parts of its range. In light of this observation, we used an ABC framework incorporating extensive simulation to explore the population history of C. *melanopterus*, focusing particularly on signals in the data that might reflect recent changes in population size. In doing so, we also use this study system as a test case to explore the extent to which metapopulation structure can hinder the detection of a recent bottleneck and propose an empirical approach and sampling scheme that addresses this issue.

## Results

### Genetic diversity and data summary

The demographic history of the metapopulation of C. *melanopterus* was examined using two datasets: the first based on estimates from a “single deme” (SID) in Northern Australia, the second based on a collection of samples taken from various locations in the Indo-Pacific, which we refer to as the “scatter” sample (SCD) (see Fig. 1, Table 1 and Supplementary Table 1). We sequenced 18 individuals in total with an average coverage of 75×. After applying strict filters and removing duplicate reads (see SI), we obtained sequence data for 995 loci for SCD and 998 loci for SID, spanning 606,647 bp and 632,160 bp respectively (Table 1 and Supplementary Table S2). The average length of the regions is ~600bp and details of the length distribution for SCD and SID are shown in Supplementary Fig. 1. Overall, 2605 and 1946 high quality single nucleotide polymorphisms (SNPs) were called for the scatter and single deme dataset, respectively. As expected, we detected a higher number of SNPs from SCD (considering samples from different locations) than SID, and SNP density is higher in introns than in exons for both datasets (no significant difference was found between SNP density in intron 5’ and intron 3’; Supplementary Table S2). We also performed a Principal Component Analysis (PCA) including all 18 samples to assess the level of population structure within our dataset (Supplementary Fig. S2). At least three distinctive clusters are suggested by the first two principal components, explaining ~40% of the variance and separating Australia from the Indo-Pacific and Oman samples. Additional population sub-structure within the Australian and the Indonesian clusters is highlighted by the third principal component (Supplementary Fig. S2, panel b and d). Overall, this confirms that the signal in our dataset is incompatible with panmixia and thus a simple demographic model of a single isolated population is not appropriate to fully describe the demographic history of these samples.

**Fig. 1:**
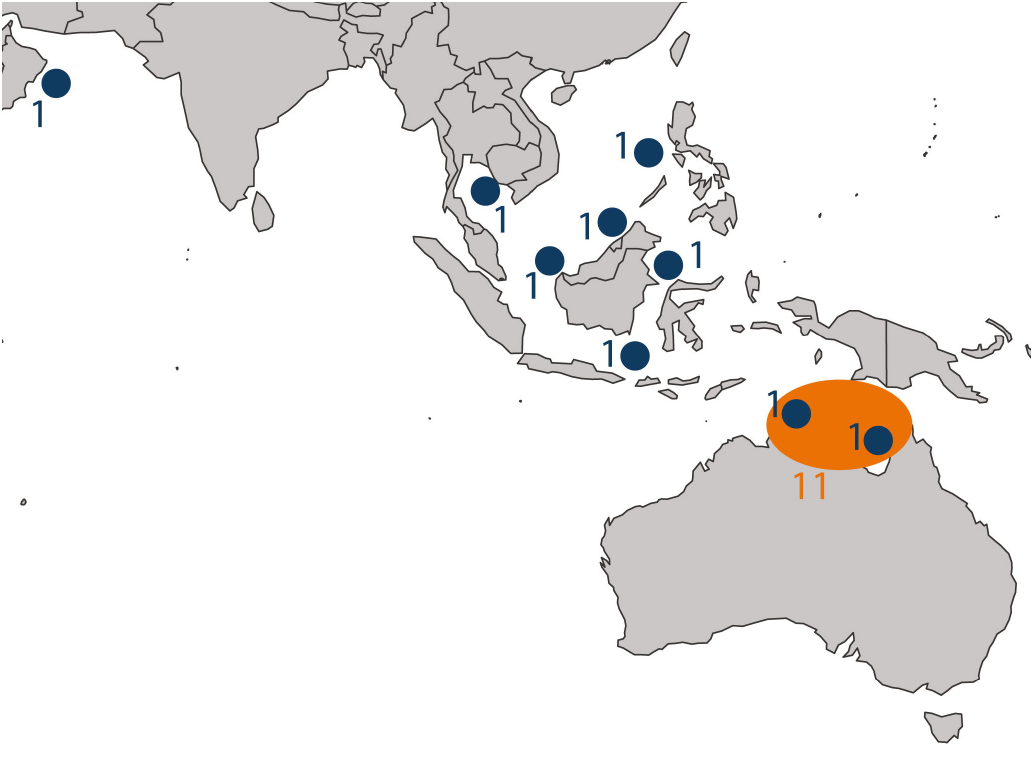
Sampling locations with sample size. Samples in blue were included in the scatter dataset (SCD, 9 samples). Samples in orange were considered for the single deme dataset (SID, 11 samples). Map of sampling locations was generated with the library “rworldmap” with R software (v3.0.2, https://cran.r-project.org/)^54^.

**Table 1:** summary of the “scatter” (SCD) and “single deme” (SID) dataset.

| Dataset | No. of samples | No. of regions | Bp sequenced | No. of SNPs | SNP density region (kb) |
| --- | --- | --- | --- | --- | --- |
| SCD | 9 | 995 | 606,647 | 2605 | 4.3 |
| SID | 11 | 998 | 632,160 | 1946 | 3.1 |

### Demographic inferences: indirect approach

First, simple population genetics models were applied to indirectly estimate the demographic parameters of the metapopulation by contrasting the characteristics of the genealogies under the two sampling schemes, SID and SCD. A constant-size model (model COS) and a demographic-change model (model CHG1) were initially compared (Fig. 2, panel a and b). Both models were found to have equal posterior probability (0.5) in SID, suggesting a pattern compatible with either a weak signal of expansion (N_mod_ was slightly higher than N_anc_, Supplementary Table S3) or a constant population size. This is consistent with the ancestral versus modern effective population size ratio (resize) computed under the CHG1 model (median = 0.48, 95% credible interval (CI) ranging from 0.03 to 2.56). In contrast to these results, model CHG1 showed a posterior probability of 0.89 when tested in SCD and a 95% CI of the resize between 0.03 and 0.54, suggesting a strong signal of expansion. This was confirmed when the effective population size was estimated at different time points in the past with an ABC-skyline reconstruction: a sharp signal of expansion was detected around ~90,000 generations before present, while a constant-size population was clearly observed in SID (Fig. 3). Under CHG1 we were able to detect the time of the expansion of the metapopulation but more recent changes in population size cannot be excluded *a priori*. Therefore, a model with two demographic changes (model CHG2; Fig. 2, panel c) was also tested in order to account for possible more recent demographic events (e.g. post-glacial expansion), as described for other marine species^30,31^. Posterior probabilities of model CHG1 and model CHG2 are similar for both datasets (Supplementary Table S4). When the effective population size was estimated at different time points in the past, CHG1 and CHG2 showed the same pattern (Fig. 3 and Supplementary Fig. S3). The estimation of T_c1_ in CHG2 for both datasets reflects the estimate of T_c_ in CHG1 suggesting that CHG2 identifies only the change in effective population size corresponding to the demographic expansion that was also identified by CHG1

**Fig. 2:**
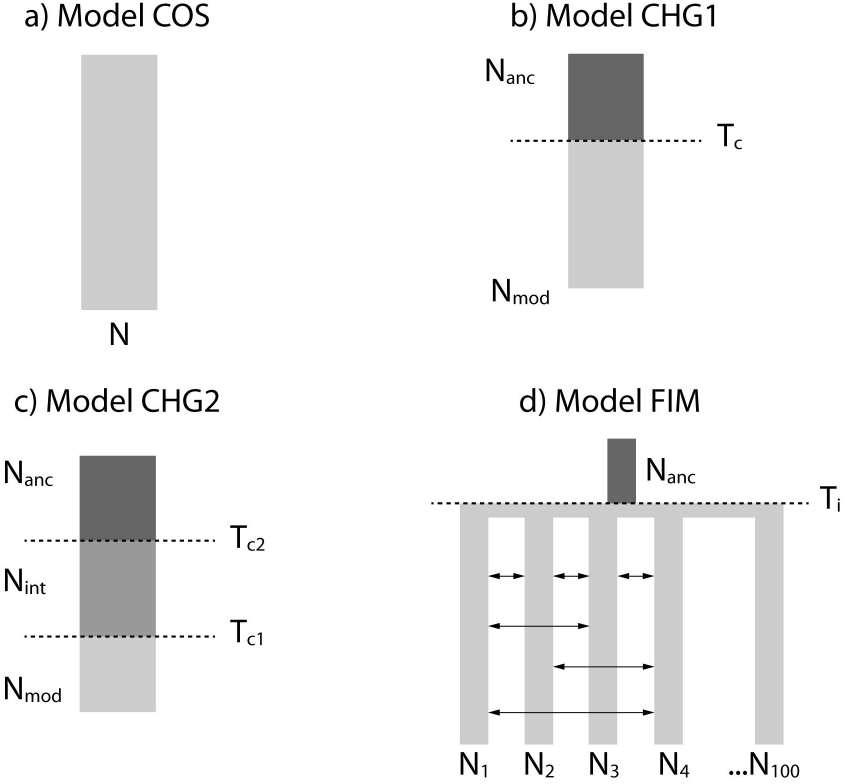
Demographic models tested for both sampling schemes. a) COS, constant-size model; b) CHG1, demographic-change model (one demographic change); c) CHG2, demographic-change model (two demographic changes); d) FIM, non-equilibrium finite island model. Nmod: modern effective population size; Nanc: ancestral effective population size; Tc, Tc1 and Tc2: time of the demographic change (in generations); Ti: time of the onset of the island (in generations).

**Fig. 3:**
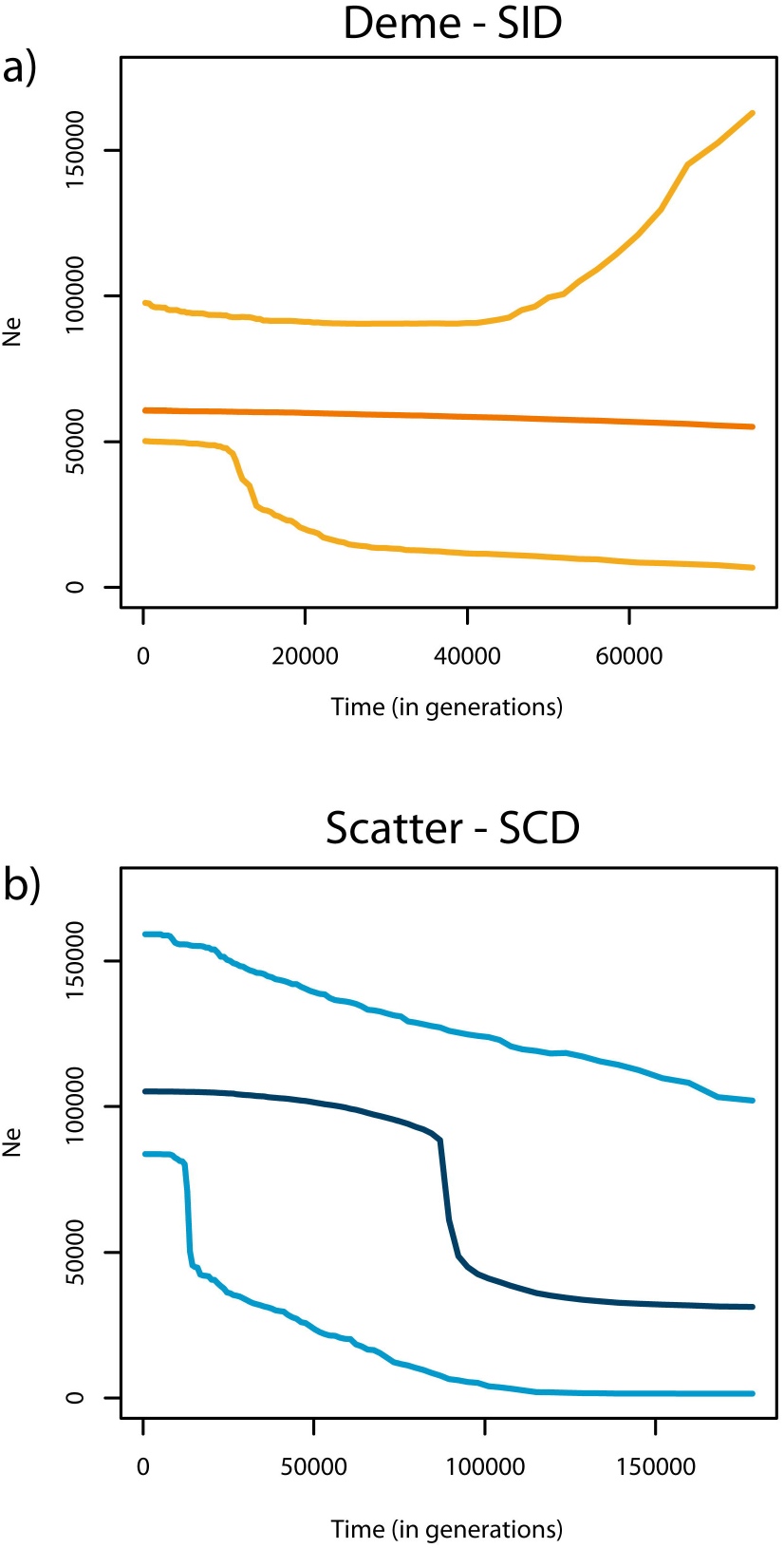
Skyline reconstruction of the effective population size through time. a) single deme (SID, in orange); b) scatter (SCD, in blue). Both skylines were reconstructed up to the mode of the mean TMRCA across loci. Median values are shown (darker lines) with the 95% high posterior density interval (lighter lines).

### Demographic inferences: direct approach

Considering the results from the indirect approach, we therefore explored the demographic signals in the data by fitting a non-equilibrium finite island model ^20^ (model FIM; Fig. 2, panel d) with 100 demes to both datasets. The choice of an island model is justified by the absence of panmixia in the PCA and by the near-uniform Gst matrix (Supplementary Table S5), suggesting similar genetic distances across all samples except Oman. Model FIM is defined by three parameters, namely *Nm*, the time of the onset of the island T_i_ and the effective size of the ancestral deme, N_anc_. Demographic parameters of the metapopulation were estimated directly for the two different sampling schemes, SCD and SID. The FIM model showed the highest posterior probability in the model choice for both SID and SCD (Table 2). Cross-validation of the model choice confirmed that FIM is distinguished from COS and CHG1 with high probability for both datasets (see SI and Supplementary Table S6). Importantly, the two datasets produced similar estimates of both *Nm* and T_i_ (Table 3). We further confirmed this result by applying FIM to the complete dataset of 18 samples (SCD+SID without shared samples) (Table 3). With this value of *Nm* ≈ 40 in the metapopulation, the signature of the ancestral expansion at T_i_ is lost in SID but not in SCD, consistent with expectation based on results in simulation studies of range expansion (see ^19,32,33^). Indeed, for similar *Nm* values the gene genealogy of lineages sampled from a single deme is a mixture of recent coalescent events occurring during the *scattering* phase and more ancient coalescent events occurring during the *collecting* phase. This generates a gene genealogy similar to that typical of a constant size population. Conversely, a scatter sample will show a signature of expansion as most of the coalescent events will occur at the time of the foundation of the metapopulation. Cross-validation of *Nm* suggests that the estimation is reliable and robust, with bias of the mode around -0.02 and -0.01 for SCD and SID respectively (Supplementary Table 7). We estimated T_i_ to have occurred ~59,000 generations ago (the average between the point estimates for the onset of the island from SCD and SID; Table 3). We observed an overestimation of the onset of the expansion in CHG1 (as T_c_) compared to FIM (as T_i_) in SCD. Cross-validation suggests a more robust and accurate estimate of T_i_ than T_c_, with an associated bias of 0.48 and 1.154 respectively (Supplementary Table S7).

**Table 2.** models posterior probability calculated as in^58^ using the closest 25,000 simulations. SCD: scatter dataset; SID: single deme dataset.

| Dataset | COS | CHG1 | FIM |
| --- | --- | --- | --- |
| SCD | 0.00 | 0.40 | 0.60 |
| SID | 0.24 | 0.31 | 0.45 |

### Detection of a recent bottleneck

We used ABC to test whether the absence of a bottleneck in our dataset reflects the real demographic history of C. *melanopterus*, or a lack of power to detect it. To this end, we simulated pseudo observed data sets (*pods*) under a modified FIM model (FIM-BOTT) characterised by two additional parameters (Supplementary Fig. S4, panel b): T_bott_ (the time when an instantaneous decrease in *Nm* occurred) and I_bott_ (the ratio of the ancestral *Nm* to the new one). When I_bott_=1 FIM-BOTT reduces to FIM (Fig. 2 panel d). FIM-BOTT is defined by two major events: the onset of the island at T_i_ and the reduction in connectivity (i.e., a reduction in the effective population size of a deme, a reduction in the migration rate, or both) at T_bott_. We simulated 1,000 *pods* under model FIM-BOTT for each parameter combination and we reanalysed them using the CHG2 model. CHG2 allows for two changes in effective population size through time and it could therefore recover both events of FIM-BOTT. We mimicked both the SID and the SCD sampling schemes and we tested twelve combinations of I_bott_ and T_bott_, namely two intensities of *Nm* reduction (100× and 1000×) at six time points (10, 50, 100, 200 500, 1000 generations ago). In this way, we were able to examine their respective behaviours when a recent bottleneck (here represented as a decrease in *Nm*) was simulated in order to obtain insight regarding the optimal sampling strategy to detect it. We plotted for comparison the results obtained by analysing *pods* simulated under FIM. For T_bott_ ≤50 generations no signal of the bottleneck was detected for either sampling scheme at any I_bott_ (Fig. 4, Supplementary Fig. S5 and S6). With progressively older T_bott_, we observed a general reduction of the estimated Ne when compared to the one estimated from pods simulated under FIM (Fig. 4, Supplementary Fig. S5, S6). This reduction was detected first in SID and only later (~500 generations ago) in SCD (Supplementary Fig. S5 and S6). However, a change in Ne through time (expected under a bottleneck) was never observed for any combination of I_bott_ and T_bott_ in SCD (Fig. 4 and Supplementary Fig. 5 and 6). SID was more affected by the change in connectivity, but a signature of a bottleneck appeared only for I_bott_=1000× and T_bott_ ≥200 (Supplementary Fig.S6). The capacity to detect a bottleneck can depend both on the real properties of the gene genealogy and on the inferential method used to analyse the data (the ABC skyline in this case). Therefore, we tested the power of our ABC-skyline approach with *pods* simulated under an unstructured model with bottleneck intensities that were comparable to those simulated in the metapopulation scenario. Specifically, we performed simulations under model CHG1-BOTT (Supplementary Fig. S4, panel a), an unstructured population with demography estimated under CHG1 (in our SCD sample) and the same combinations of I_bott_ and T_bott_ as above. Here, the parameter I_bott_ refers to the decrease in Ne, while for FIM-BOTT it referred to the decrease in *Nm*. Ne estimated from *pods* simulated under CHG1-BOTT drops compared to the values estimated from *pods* simulated under CHG1 as recently as 10 generations ago (Fig. 4). Moreover, a clear signature of a bottleneck appears as soon as T_bott_=50 (Fig. 4).

**Fig. 4:**
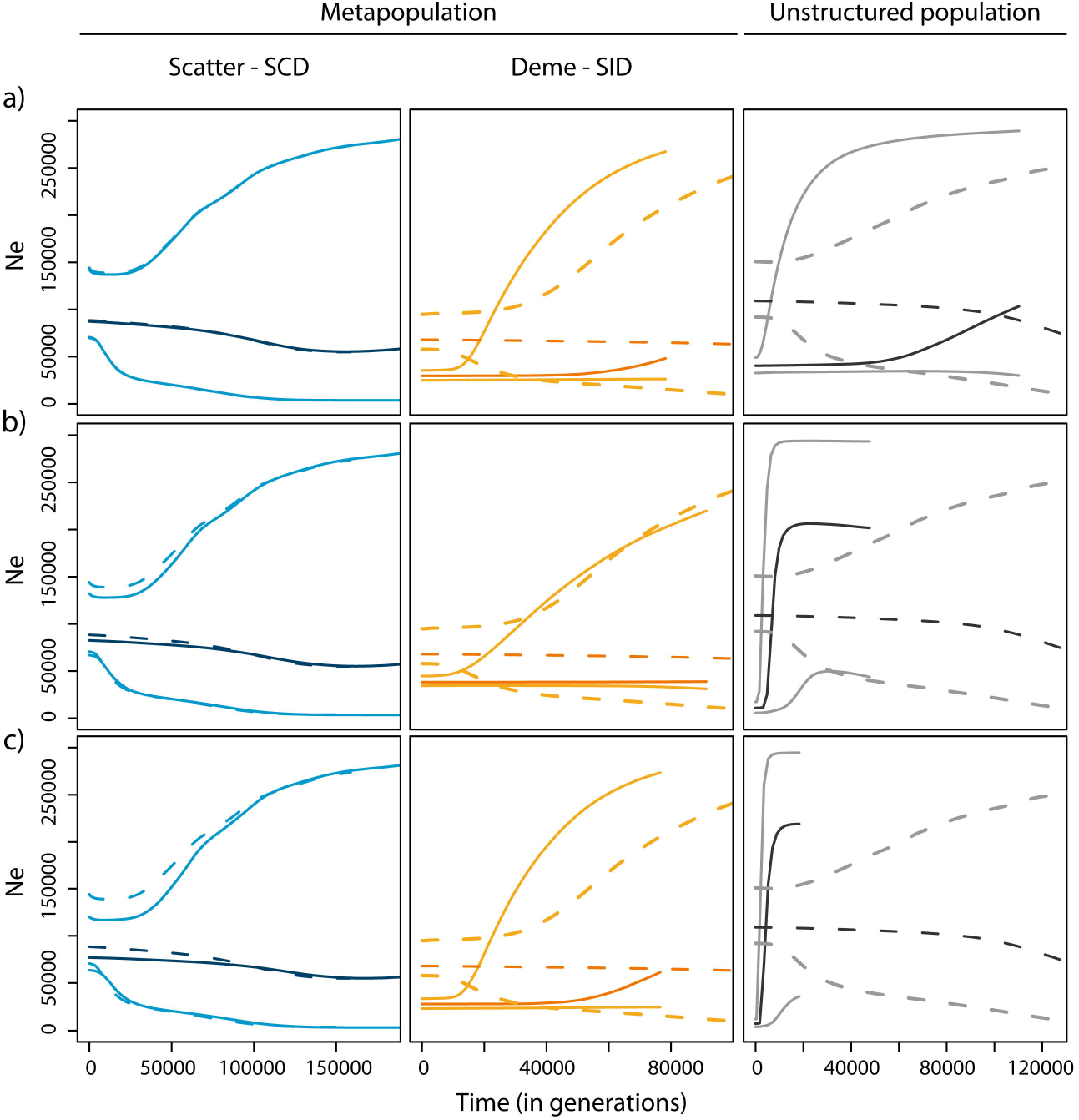
Average skyline reconstruction under model CHG2 of pseudo-observed data sets (*pods*) simulated with model FIM and FIM-BOTT in a metapopulation (for SCD and SID) and in an unstructured population (model CHG1 and CHG1-BOTT). Data were simulated with I_bott_=1000 at: a) T_bott_=10 generations; b) T_bott_=50 generations; c) T_bott_=100 generations before present. An average skyline reconstruction is shown across 1000 simulations. Solid lines: scenario with bottleneck (model FIM-BOTT and CHG1-BOTT); dashed line: scenario without bottleneck (model FIM and CHG1). Median values are shown (darker lines) with the 95% high posterior density interval (lighter lines).

Similar to the reasoning followed when studying the real data, we further analysed the *pods* simulated under FIM-BOTT for all combinations of I_bott_, T_bott_ and sampling scheme with the FIM model. We plotted the median of the posterior distributions of *Nm* estimated from each *pods* (hereafter, *Nm_est_*). To quantify the reduction of *Nm_est_* due to the simulated decrease in *Nm*, we show as comparison *Nm_est_* obtained from *pods* simulated under FIM (Fig. 5, Supplementary Fig. S7). First of all, we note that both sampling schemes can correctly recover *Nm* when *pods* are simulated with the FIM model, indicating that this is an appropriate inferential procedure (Fig. 5). We then observed that the two sampling scheme respond differently when *pods* are simulated under FIM-BOTT. Generally, T_bott_ seems to have a stronger effect on *Nm_est_* than does I_bott_ (Fig. 5, Supplementary Fig. S7). SCD does not show any significant reduction in *Nm_est_* until 500 generations ago, while SID starts showing a decrease almost immediately, after just 10 generations. The decline of *Nm_est_* in SID is faster when I_bott_=1,000 than for I_bott_=100. In both cases, the distributions of *Nm_est_* for SID and SCD up to 500 generations ago show a significant difference. At 1000 generations, both datasets show the same pattern with severe decreases in *Nm_est_* (Fig. 5, Supplementary Fig. S7).

**Fig. 5:**
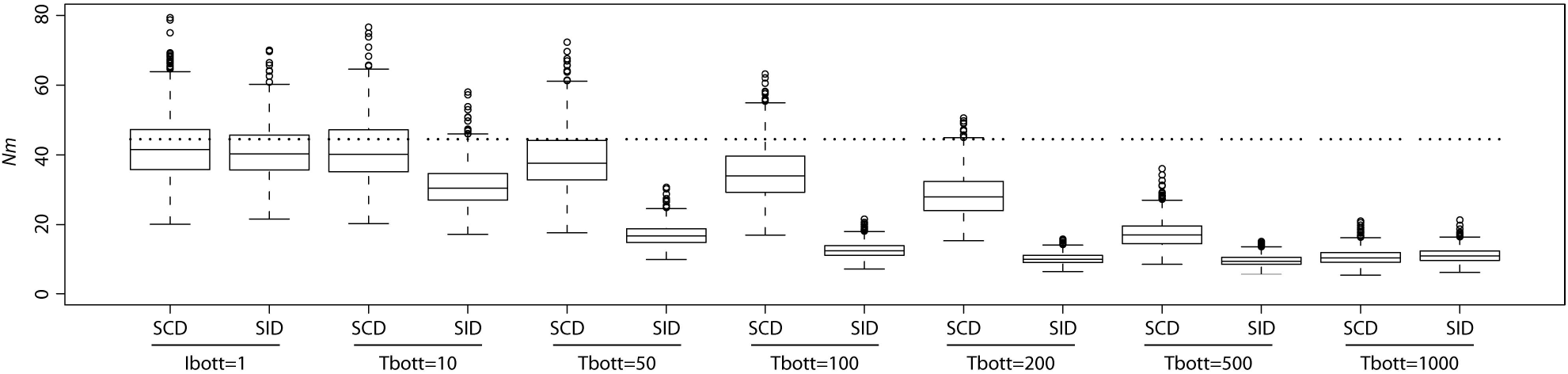
Distribution of the median of *Nm* estimated under model FIM from pseudo observed data sets (*pods*) generated with FIM-BOTT and FIM. Data were simulated for both sampling schemes with I_bott_=100 and various T_bott_(10, 50, 100, 200, 500 and 1000 generations ago). Dotted lines represent the value of *Nm* used to simulate pods under FIM model.

## Discussion

In this study we present a novel approach to population genomics that is suitable for application to non-model organisms, particularly those for which it is difficult to obtain large sample sizes. The novelty stems from both the technology applied to produce NGS sequence data and the sampling scheme adopted for the population genomics inferences. First, we used a recently developed target gene capture approach paired with NGS ^23^ that, so far, has only been used for phylogenomics inference34. Second, we exploit intrinsic differences in the shape of gene genealogies that result from the *collecting* and *scattering* phases of metapopulations^16,18,32^ to demonstrate that it is possible to make demographic inferences indirectly based on a few carefully chosen individuals when using a sufficiently large number of loci. We chose to test the utility of our framework using the black tip reef shark, C. *melanopterus*, because we felt that this species was a good example of the practical challenges that are faced by population and conservation geneticists that are working with non-model systems. Firstly, individuals of this species have a small home range and exhibit strong site fidelity, making it a good example of a non-model organism that exhibits metapopulation structure. Indeed, this species has previously been shown to exhibit extensive population structure and restricted movements in French Polynesia ^35-37^, the Great Barrier Reef ^38^ and Western Australia39. Secondly, they mostly occur in remote regions of the world and, as is the case for the majority of free ranging marine animals, it is logistically difficult to sample them in large numbers. Finally, the International Union for the Conservation of Nature has listed C. *melanopterus* as “near threatened”, meaning that any information that can be gleaned regarding the demography and structure of this species can have implications for management. These attributes make C. *melanopterus* a well-suited system for to test the efficacy of our approach for reconstructing the evolutionary dynamics, historical demography and associated conservation implications in a non-model species that has a high degree of metapopulation sub-structure, but from which widespread sampling is difficult.

Our approach derives its effectiveness from the separation of time scales (i.e., *scattering* vs *collecting* phases), first described elegantly by Wakeley (1999), for the gene genealogies in an island model and later shown to be valid for many metapopulation models, i.e., stepping stone ^18^, spatially continuous ^40^ and range expansion ^19^. Our indirect approach based on the comparison of the genealogical history at the two sampling levels appears robust to mis-specification of the metapopulation model, but it cannot provide precise estimates of the demographic parameters. To further characterise the underlying demography, we analysed our data directly using a non-equilibrium finite island model. While other metapopulation models might better fit the data, they come at the risk of overparameterisation (i.e., non-symmetric island model, spatial models as a stepping stone). Herein, we demonstrate statistically that even a relatively simple model such as the FIM provides a better proxy to the data and more realistic description of a species history than models that ignore population structure (Table 2).

The relative length of the *scattering* vs *collecting* phase determines the shape of the gene genealogy and is mostly influenced by *Nm*^16,19^. This holds true for lineages sampled within a deme (our SID) and lineages scattered throughout the range of the species (our SCD). If an expansion occurred in the species through the colonisation of new habitat, high *Nm* values will determine a signature of population growth at all sampling levels 18. For decreasing *Nm*, the signature of expansion is first lost when sampling lineages from a single deme and then in a scattered sample. We found a significant signature of expansion in SCD and a constant population size in SID. Ray et al. (2003) suggested that at *Nm* < 50 the genealogy in a single deme would mostly resemble that of a constant size population (or even of a population reducing in size) in a range expansion model. When we fit the FIM model independently to both SCD and SID, we found a metapopulation characterised by a value of *Nm* ~40 (95% CI: 20-110), with an onset of formation around ~59,000 (95% CI: 6,000-65,000) generations ago. This level of connectivity is consistent with philopatry in this species, as previously described in Moorea and Tetiaroa (French Polynesia)^35^. Unfortunately our sampling scheme does not allow us to test males and females separately, but it would be interesting to investigate possible differences in the level of connectivity related to sex-specific migration. We further apply the same FIM model to the whole dataset and we obtained similar values of both *Nm* and T_i_ (Table 3). In summary, the indirect approach of comparing gene genealogies under different sampling schemes and directly applying the FIM model (applied to three largely independent datasets) provided consistent results, suggesting that this approach is effective for reconstructing the demography of this species. We stress that even though there is no spatial component in the FIM model, the time of formation of the metapopulation is analogous to the expansion time in a range expansion model. Therefore, we speculate that this time estimate probably represents the colonisation of the Indian Ocean by C. *melanopterus*, through a range expansion. Vignaud et al.^29^ found evidence for genetic structure and isolation by distance at a larger geographic scale (i.e., Pacific and Indian Ocean) for this species. Clearly, a FIM model would not be a good approximation in this case and a non-equilibrium stepping-stone or a range expansion model might be more appropriate. However, our individuals (with the exception of Oman) come from a restricted geographic area so that the spatial component of the demographic model can be neglected. Indeed, we simulated from the posterior distribution of the FIM the expected *Fst* between two demes at 14 independent microsatellites loci with the same mutation rate as in Vignaud et al.^29^. We found the *Fst* distribution to be compatible with the value they found between East and West Australia, which are more distant than the samples from our study area. This further suggests that our model is appropriate for our specific dataset, but we are aware that it might not provide an accurate description of the C. *melanopterus* demography over its whole range.

We made some simplifying assumptions when analysing the C. *melanopterus* data. We used a single step change of effective population size in our indirect approach and a constant *Nm* when fitting the FIM model. Obviously, the real evolutionary histories of species are likely to be more complex and it may be challenging to distinguish between local events (such as a bottleneck or expansion in a deme) and changes in connectivity/number of demes in metapopulations ^16,41^. Climate change and anthropogenic activity are reported to have impacted several marine species, including some sharks ^30,42-44^. A recent bottleneck in C. *melanopterus* was identified in Moorea and attributed to human activity ^29^. In an effort to distinguish between patterns that might be caused by metapopulation structure from those caused by localized bottlenecks we subjected the data to analysis using both SID and SCD. We did not find any evidence for a more recent event than the metapopulation expansion of C. *melanopterus* (Supplementary Fig. S3), ruling out the possibility of a demographic change related to the post-glacial maximum (~20-15 KYA) as has been proposed for other sharks 30. The distribution of C. *melanopterus* is confined to reef flats and sheltered lagoons and the effects of the last glacial maximum appear to have had little impact on this species. However, recent and mild bottlenecks are more challenging to detect using coalescent analyses ^45,46^. Indeed, many marine species do not show clear signatures of a bottleneck despite being threatened by overfishing for many generations ^47,48^. Nevertheless, meta-analysis studies have shown general reduction in diversity in overfished species49. These issues motivated us to investigate how best to detect a recent reduction in effective size in a metapopulation. To this end, we performed a simulation study focusing on a metapopulation with the same demography as that estimated in our C. *melanopterus* data but experiencing a recent bottleneck. Obviously, a more extensive investigation of parameter space would be interesting but is beyond the scope of this paper.

**Table 3.** parameter estimation under the finite island model (FIM). N_anc_: effective population size of the ancestral deme; T_i_: time of the onset of the island (in generations); *Nm*: product of the effective population size N and the migration rate *m* for each deme; SCD: scatter dataset (9 individuals); SID: single deme dataset (11 individuals); pooled dataset: SCD+SID without shared samples (18 individuals).

| Model FIM | Median | Mode | 0.025<sup>a</sup> | 0.975<sup>a</sup> | Prior<sup>b</sup> |
| --- | --- | --- | --- | --- | --- |
| **SCD** |  |  |  |  |  |
| $N_{anc}$ | 33,567 | 42,704 | 1604 | 62,242 | U: 100-100,000 |
| $T_i$ | 72,760 | 56,354 | 24,124 | 141,628 | U: 100-200,000 |
| $Nm$ | 52.85 | 47.17 | 38.80 | 109.77 | U: 0.05-250 |
| **SID** |  |  |  |  |  |
| $N_{anc}$ | 31,016 | 30,737 | 1571 | 86,026 | U: 100-100,000 |
| $T_i$ | 63,431 | 61,671 | 6095 | 163,634 | U: 100-200,000 |
| $Nm$ | 42.04 | 38.61 | 28.51 | 117.80 | U: 0.05-250 |
| **Pooled dataset** |  |  |  |  |  |
| $N_{anc}$ | 52,541 | 58,148 | 8176 | 84,492 | U: 100-100,000 |
| $T_i$ | 65,004 | 18,329 | 4255 | 176,692 | U: 100-200,000 |
| $Nm$ | 58.7 | 51.4 | 38.3 | 150.9 | U: 0.05-250 |

<sup>a</sup>Upper and lower limits of the 95% credible interval about the estimated mode.
<sup>b</sup>U, uniform probability, in the range of the two values.

This simulation study had three goals: i) to better understand the recent demographic history of C. *melanopterus*; ii) to characterise the behaviour of metapopulations experiencing recent bottlenecks; iii) to evaluate the relative merits of our different sampling schemes. Analyses of the simulated data paralleled those adopted for the real data, i.e., using the direct and indirect approaches. The indirect approach revealed two important features: i) SCD fails to detect evidence of a bottleneck, even at high intensities, showing a pattern that is similar to the no-bottleneck case (Fig. 4); ii) SID similarly does not show evidence of bottleneck, but it shows a reduction in genetic diversity compared to the no-bottleneck case at both intensities (100× and 1000×), within just 10 generations. The genetic consequence of the bottleneck on a SID sample is the reduction in the estimated effective size compared to the no-bottleneck scenario (Fig. 4, Supplementary Fig. S5 and S6). However, we could not have detected such a decrease of the effective size without some baseline reference values, suggesting that it is difficult to assess whether a species is critically endangered when it is part of a structured metapopulation. This might explain why many overfished species do not show a genetic signature of bottleneck despite being threatened.

The direct approach confirmed that the two sampling schemes behave differently after a bottleneck. We computed *Nm* using the FIM model and we found, as expected, that a stable metapopulation shows a similar estimate of *Nm* under both sampling schemes (Fig. 5 and Supplementary Fig. S7). Conversely, a decrease in the *Nm* estimated from SID, but not from SCD, is observed under a bottleneck scenario (Fig. 5 and Supplementary Fig. S7), suggesting that the comparison of the two datasets may help detecting recent changes in effective population size. These results (both the ABC-skyline models and the *Nm* estimates) can be interpreted from a coalescent point of view: a recent bottleneck will produce a burst of coalescent events only at the deme level (i.e., during the *scattering* phase), having no impact at all on the *collecting* phase even for high intensity of reduction. A scattered sample, which does not have the *scattering* phase, will not be affected by recent bottleneck events while a local single deme sample will. These findings suggest that metapopulations are more resilient than unstructured populations to recent changes in effective size (or connectivity), implying that taking population structure into account is crucial when inferring the demography of a species. Most species are structured, including those that are endangered. In conservation studies we need to detect changes in effective population size on the order of tens to, at most, hundreds of generations. Our method can be therefore be usefully applied in a conservation setting as it is able to detect such recent changes. We expect that it can be easily applied to both critically endangered (e.g. the daggernose shark) and commercially exploited (e.g.. the Atlantic cod) species to better design appropriate conservation management plans. We now envisage exploring the power of our approach under a wider spectrum of parameters in order to provide guidelines for conservation studies, but this is beyond the scope of this work. In the case of the C. *melanopterus* data examined in the current study, the single deme sampled from Northern Australia has not experienced a recent bottleneck. However, we cannot exclude the possibility that this may have occurred for the other demes of the metapopulation, where conditions may have been more extreme than those simulated here.

In summary, we present a recently developed target gene capture approach used here for the first time to perform inferences at the intra-specific level. By exploiting theoretical and simulation results on the coalescent history of metapopulations, we show how few carefully collected specimens can provide information about the demography of a species when many loci are available. This strategy can minimise sampling effort and associated cost of data production, making population genomic analysis of non-model organisms more feasible. We showed that this approach allows refined estimates of connectivity among demes relative to traditional population genetics approaches and that the sampling scheme adopted is particularly effective when investigating recent changes in effective population size and, potentially, migration rates. This opens new avenues for theoretical developments on the conservation management and recovery of endangered species.

## Materials and Methods

### Sampling, library preparation and sequencing

A total of 18 samples of C. *melanopterus* were included from 8 different locations (Fig. 1): Northern Territory and Queensland, Australia (N=11), Oman (N=1), Philippines (N=1), Thailand (N=1), Malaysia (N=1), South Kalimantan, Indonesia (N=1), West Kalimantan, Indonesian Borneo (N=1) and East Kalimantan, Indonesia (N=1). Samples were divided in two datasets called “scatter” (SCD) and “single deme” (SID) with 9 and 11 samples respectively (Table 1, Supplementary Table S1). The SCD dataset aims to mimic a “scatter sample”, sensu Wakeley, where all samples come from independent geographical locations distributed in close proximity to each other. We also included one sample from Oman in order to infer the demographic history of the metapopulation at a larger scale and to cover a more extensive area of the Indian Ocean. Muscle tissue was extracted from dead specimens collected from local fish markets and then stored in 95% ethanol. Genomic DNA was extracted using the E.Z.N.A Tissue DNA Kit (Omega Bio-Tek, Inc Norcross, GA) according to the manufacturer’s instructions. Illumina sequencing libraries (500bp) were prepared and amplified by PCR prior to two rounds of target gene capture (targeting 1077 autosomal regions) using a ‘touchdown’ DNA hybridization approach^23^. Resulting enriched libraries were re-amplified to incorporate a sample specific index, pooled and sequenced paired-end (250 bp reads) on an Illumina MiSeq benchtop sequencer (Illumina, Inc, San Diego, CA). Sequence reads associated with each sample were identified and sorted by their respective indices^23^.

### De novo reference sequence assembly

Briefly, sequence read data from several individuals was used to build a haploid reference sequence for the 1077 target exons and associated introns. Adapters and low quality reads were removed from the raw sequence read data and contigs were assembled *de novo*. On target material was identified by determining the orthology of assembled contigs to a set of core orthologs identified across model vertebrates (see SI for details). For each exon and intron (5’ and 3’) the longest contig across all sequenced individuals was retained and used to assemble a reference sequence that is 1,293,710 bp including exons and introns (5’ and 3’).

### Read mapping, variant calling and filtering

A Burrows-Wheeler Aligner (BWA) ^50^ was used to map reads to the reference sequence generated by de novo assembly ^51^. Duplicate read marking was performed with Picard v.1.129 (http://broadinstitute.github.io/picard/), local realignment with the Genome Analysis ToolKit (GATK) v3.3-0 ^52^ and genotype calling with samtools v1.1 ^53^. Raw variants were filtered using custom Perl (v5.18.2) and R (3.0.2) scripts (Supplementary Table S8). Filtering was based on strand bias, minimum depth 6×, removal of triallelic sites, removal of sites with heterozygous calls in more than 80% of samples and removal of any missing calls in order to have a dataset without any missing genotypes, reducing background noise and uncertainty (see SI for details).

### Descriptive analyses

Principal component analysis (PCA) was performed with the function “prcomp” in the R environment ^54^. A Gst distance matrix^55^ was calculated with Arlequin 3.5^56^ (see SI for details).

### Demographic Inference

We developed an ABC ^24^ framework to estimate parameters and compare demographic models. The folded site frequency spectrum (SFS) and total number of SNPs were used as summary statistics, to avoid phasing issues. Mutation and recombination rates were allowed to vary across loci using hyperprior distributions (see SI for details). Generation time was set to seven years ^36^. Four demographic models were tested for both datasets (Fig. 2). Model COS represents a constant-size population model described only by the effective population size (N) (Fig. 2, panel a). Model CHG1 represents a single instantaneous demographic-change model occurring at the time T_c_ (Fig. 2, panel b). N_anc_ and N_mod_ are the ancestral and modern effective population sizes respectively. Values of the ancestral versus modern effective population size ratio (resize, N_anc_/N_mod_) greater than 1 are compatible with a reduction in effective population size while values lower than 1 suggest a population expansion. If this ratio is equal to 1, the scenario is compatible with a constant-size population. Model CHG2 represents a demographic-change model in which two events, occurring at times T_c1_ and T_c2_, cause changes in population size described by three effective population sizes (N_anc_, N_int_ and N_mod_ for ancestral, intermediate and modern respectively) (Fig. 2, panel c). Model FIM represents a non-equilibrium finite island model with 100 demes (N_1_…N_100_) described by *Nm* as the product between the effective population size (N) and the migration rate (m) (Fig. 2, panel d). N_anc_ is the effective size of the ancestral deme from which the island originated at the time T_i_. One hundred demes was chosen in order to approximate the large number of demes that are necessary to describe the coalescent history in metapopulation models in terms of the *scattering* and *collecting* phases^20^.

We generated 100,000 simulations for each demographic model using fastsimcoal2 v2.5.1^57^. Each simulation includes 995 and 998 gene genealogies for SCD and SID respectively, to be consistent with our real data. Prior distributions are displayed in Table 3, Supplementary Table S3 and S9. The posterior probabilities of each model were calculated by a weighted multinomial logistic regression^58^ for which we retained the best 25,000 simulations. The parameters of the best-fitting model were estimated from the 5,000 simulations closest to the observed dataset using a local linear regression according to ^24^. Posterior distributions for model CHG1 and FIM are shown in Supplementary Fig. 8 and 9.

Both CHG1 and CHG2 models were used to graphically reconstruct the variation of effective population size through time. To this end, for each combination of parameters retained by the ABC algorithm (5,000 in our case), we recorded the effective size at specific time points. The median value of the posterior distribution of the effective size at each time point was calculated together with the 95% credible interval and plotted against time to obtain an ABC-skyline reconstruction (Fig. 3, 4, Supplementary Fig. S3, S5 and S6). Time points were defined by randomly extracting values from an exponential distribution with a rate calibrated with the upper bound (97.5%) of the estimated mean TMRCA across loci. In this way all generated time points were not greater than 97.5 % of the estimated mean TMRCA across loci. Moreover, recent time points (0, 25, 50, 100, 200, 300, 400 and 500 generations ago) were manually added to increase the resolution towards recent events. Analyses were performed in the R environment^54^ with the library abc ^59^.

We performed cross-validation for both model selection and parameter estimation by randomly generating *pods* from the prior distributions under each model. For each cross-validation experiment we generated 1000 *pods* and we applied the same inferential procedure as for the observed data (see SI, Supplementary Fig. S10 and S11). In addition, a posterior predictive test ^60^ was carried out to test whether the data can be reproduced under a specific demographic model. Bayesian p-values computed from the posterior distribution of the number of polymorphic sites showed that none of the four models in both datasets could be rejected (SI, Supplementary Fig. S12).

### Recent Bottleneck

We tested for signatures of a recent population bottleneck in the metapopulation data set for both sampling schemes. We restricted our ABC analyses to a metapopulation with demographic parameters consistent with those estimated from the empirical data examined in this study. We simulated *pods* under a modified FIM model (FIM-BOTT) characterised by two additional parameters (Supplementary Fig. S4, panel b): T_bott_ (the time when an instantaneous decrease in *Nm* occurred) and I_bott_ (the ratio of the ancestral *Nm* to the new one). When I_bott_=1 FIM-BOTT reduces to FIM (Fig. 2, panel d). We tested twelve combinations of parameters, namely two I_bott_ (100× and 1000×) and six T_bott_ (10, 50, 100, 200, 500 and 1000 generations ago). For each combination of parameters, we simulated 1000 *pods* for both SCD and SID (Supplementary Fig. S4, panel b). To further understand the effect of metapopulation structure on bottlenecks, we performed additional simulations of an unstructured population under a modified CHG1 model (CHG1-BOTT, Supplementary Fig. S4, panel a). In this case, *pods* were simulated with demographic parameters estimated in our real SCD data under model CHG1, to which we added a recent bottleneck characterised by I_bott_ and T_bott_. Note that here I_bott_ represents the ratio between the ancestral effective population size *Ne* and the new one (while in FIM-BOTT it represents the ratio of the ancestral *Nm* to the new one). We tested the same twelve parameters combinations of I_bott_ and T_bott_ that we previously used when simulating FIM-BOTT. CHG1-BOTT reduces to CHG1 when I_bott_ =1. All *pods* simulated under both FIM-BOTT (for both sampling schemes) and CHG1-BOTT were analysed using the ABC-skyline produced by the CHG2 model (Fig. 2, panel c). We applied the same settings used for the real data and for each combination of parameters we plotted the average of the *Ne* through time across the 1,000 *pods* (see SI). All bottleneck scenarios were compared to the ABC-skyline reconstructed from *pods* simulated under FIM or CHG1. Finally, *pods* simulated under FIM-BOTT were also analysed using FIM to estimate *Nm*. We compared these values to those estimated from *pods* generated under FIM.

## Acknowledgments

The authors thank Giorgio Bertorelle and Oscar Lao for helpful comments and suggestions and Laurent Excoffier for sharing fastsimcoal v2.5.2 before publication. Data analysis was performed on the computer cluster Genotoul bioinformatics platform (www.bioinfo.genotoul.fr). This work was supported by the Agence Nationale de la Recherche (ANR-12-BSV7-0012 to P.M.D.) and by the US National Science Foundation (DEB-1132229).

## Author Contributions

S.M., S.P. and G.N. designed research; M.H, C.L. and S.C. contributed with new reagents and analytical tools; P.M.D., S.C., S.M. and G.N. performed research; P.M.D. and S.M. analysed data; M.V. gave comments on the manuscript and project; P.M.D., S.C., S.M and G.N. wrote the paper.

## Data accessibility

A vcf file containing all variants used in this study is being submitted to the Dryad database (http://datadryad.org/).

## References

1 Nadachowska-Brzyska, K. et al. Demographic divergence history of pied flycatcher and collared flycatcher inferred from whole-genome re-sequencing data. PLoS Genet 9, e1003942, (2013).

2 Romiguier, J. et al. Comparative population genomics in animals uncovers the determinants of genetic diversity. Nature 515, 261–263, (2014).

3 Emerson, K. J. et al. Resolving postglacial phylogeography using high-throughput sequencing. Proc Natl Acad Sci U S A 107, 16196–16200, (2010).

4 Li, S. & Jakobsson, M. Estimating demographic parameters from large-scale population genomic data using Approximate Bayesian Computation. BMC Genet 13, 22, (2012).

5 Roux, J., Rosikiewicz, M. & Robinson-Rechavi, M. What to compare and how: Comparative transcriptomics for Evo-Devo. Journal of Experimental Zoology Part B: Molecular and Developmental Evolution, (2015).

6 Arnold, B., Corbett-Detig, R. B., Hartl, D. & Bomblies, K. RADseq underestimates diversity and introduces genealogical biases due to nonrandom haplotype sampling. Mol Ecol 22, 3179–3190, (2013).

7 Huang, H. & Knowles, L. L. Unforeseen Consequences of Excluding Missing Data from Next-Generation Sequences: Simulation Study of RAD Sequences. Syst Biol, (2014).

8 Leache, A. D. et al. Phylogenomics of Phrynosomatid Lizards: Conflicting Signals from Sequence Capture versus Restriction Site Associated DNA Sequencing. Genome Biol Evol 7, 706–719, (2015).

9 Jones, M. R. & Good, J. M. Targeted capture in evolutionary and ecological genomics. Mol Ecol, (2015).

10 Eriksson, A. & Manica, A. The doubly conditioned frequency spectrum does not distinguish between ancient population structure and hybridization. Mol Biol Evol 31, 1618–1621, (2014).

11 Peter, B. M., Wegmann, D. & Excoffier, L. Distinguishing between population bottleneck and population subdivision by a Bayesian model choice procedure. Mol Ecol 19, 4648–4660, (2010).

12 Heller, R., Chikhi, L. & Siegismund, H. R. The confounding effect of population structure on Bayesian skyline plot inferences of demographic history. PLoS One 8, e62992, (2013).

13 Chikhi, L., Sousa, V. C., Luisi, P., Goossens, B. & Beaumont, M. A. The confounding effects of population structure, genetic diversity and the sampling scheme on the detection and quantification of population size changes. Genetics 186, 983–995, (2010).

14 Currat, M. & Excoffier, L. Strong reproductive isolation between humans and Neanderthals inferred from observed patterns of introgression. Proc Natl Acad Sci U S A 108, 15129–15134, (2011).

15 Francois, O., Blum, M. G., Jakobsson, M. & Rosenberg, N. A. Demographic history of european populations of Arabidopsis thaliana. PLoS Genet 4, e1000075, (2008).

16 Wakeley, J. Nonequilibrium migration in human history. Genetics 153, 1863–1871, (1999).

17 Wakeley, J. The coalescent in an island model of population subdivision with variation among demes. Theor Popul Biol 59, 133–144, (2001).

18 Stadler, T., Haubold, B., Merino, C., Stephan, W. & Pfaffelhuber, P. The impact of sampling schemes on the site frequency spectrum in nonequilibrium subdivided populations. Genetics 182, 205–216, (2009).

19 Ray, N., Currat, M. & Excoffier, L. Intra-deme molecular diversity in spatially expanding populations. Mol Biol Evol 20, 76–86, (2003).

20 Wakeley, J. Segregating sites in Wright's island model. Theor Popul Biol 53, 166–174, (1998).

21 Papastamatiou, Y. P., Lowe, C. G., Caselle, J. E. & Friedlander, A. M. Scale-dependent effects of habitat on movements and path structure of reef sharks at a predator-dominated atoll. Ecology 90, 996–1008, (2009).

22 Papastamatiou, Y. P., Friedlander, A. M., Caselle, J. E. & Lowe, C. G. Long-term movement patterns and trophic ecology of blacktip reef sharks (*Carcharhinus melanopterus*) at Palmyra Atoll. J Exp Mar Biol Ecol 386, 94–102, (2010).

23 Li, C., Hofreiter, M., Straube, N., Corrigan, S. & Naylor, G. J. Capturing protein-coding genes across highly divergent species. Biotechniques 54, 321–326, (2013).

24 Beaumont, M. A., Zhang, W. & Balding, D. J. Approximate Bayesian computation in population genetics. Genetics 162, 2025–2035, (2002).

25 Bertorelle, G., Benazzo, A. & Mona, S. ABC as a flexible framework to estimate demography over space and time: some cons, many pros. Mol Ecol 19, 2609–2625, (2010).

26 Heupel, M. Carcharhinus melanopterus, www.iucnredlist.org (2009).

27 Field, I. C. et al. Changes in size distributions of commercially exploited sharks over 25 years in northern Australia using a Bayesian approach. Fish Res 125, 262–271, (2012).

28 Henderson, A., Al-Oufi, H. & McIlwain, J. Survey, status and utilization of the elasmobranch fishery resources of the Sultanate of Oman. (Sultan Qaboos University, Muscat, 2007).

29 Vignaud, T. M. et al. Blacktip reef sharks, Carcharhinus melanopterus, have high genetic structure and varying demographic histories in their Indo-Pacific range. Mol Ecol 23, 5193–5207, (2014).

30 Portnoy, D. S. et al. Contemporary population structure and post-glacial genetic demography in a migratory marine species, the blacknose shark, Carcharhinus acronotus. Mol Ecol 23, 5480–5495, (2014).

31 Marko, P. B. et al. The 'Expansion-Contraction' model of Pleistocene biogeography: rocky shores suffer a sea change? Mol Ecol 19, 146–169, (2010).

32 Mona, S., Ray, N., Arenas, M. & Excoffier, L. Genetic consequences of habitat fragmentation during a range expansion. Heredity (Edinb) 112, 291–299, (2014).

33 Wegmann, D., Currat, M. & Excoffier, L. Molecular diversity after a range expansion in heterogeneous environments. Genetics 174, 2009–2020, (2006).

34 Li, C. et al. DNA capture reveals transoceanic gene flow in endangered river sharks. Proc Natl Acad Sci U S A 112, 13302–13307, (2015).

35 Mourier, J. & Planes, S. Direct genetic evidence for reproductive philopatry and associated fine-scale migrations in female blacktip reef sharks (*Carcharhinus melanopterus*) in French Polynesia. Mol Ecol 22, 201–214, (2013).

36 Mourier, J., Mills, S. C. & Planes, S. Population structure, spatial distribution and life-history traits of blacktip reef sharks Carcharhinus melanopterus. J Fish Biol 82, 979–993, (2013).

37 Vignaud, T., Clua, E., Mourier, J., Maynard, J. & Planes, S. Microsatellite analyses of blacktip reef sharks (*Carcharhinus melanopterus*) in a fragmented environment show structured clusters. PLoS One 8, e61067, (2013).

38 Chin, A., Tobin, A. J., Heupel, M. R. & Simpfendorfer, C. A. Population structure and residency patterns of the blacktip reef shark Carcharhinus melanopterus in turbid coastal environments. J Fish Biol 82, 1192–1210, (2013).

39 Speed, C. W. et al. Spatial and temporal movement patterns of a multi-species coastal reef shark aggregation. Mar Ecol Prog Ser 429, 261-U618, (2011).

40 Wilkins, J. F. A separation-of-timescales approach to the coalescent in a continuous population. Genetics 168, 2227–2244, (2004).

41 Mazet, O., Rodriguez, W. & Chikhi, L. Demographic inference using genetic data from a single individual: Separating population size variation from population structure. Theor Popul Biol 104, 46–58, (2015).

42 Hernández, S. et al. Demographic history and the South Pacific dispersal barrier for school shark (Galeorhinus galeus) inferred by mitochondrial DNA and microsatellite DNA mark. Fish Res 167, 132–142, (2015).

43 Vignaud, T. M. et al. Genetic structure of populations of whale sharks among ocean basins and evidence for their historic rise and recent decline. Mol Ecol 23, 2590–2601, (2014).

44 O’Leary, S. J. et al. Genetic Diversity of White Sharks, Carcharodon carcharias, in the Northwest Atlantic and Southern Africa. Journal of Heredity, (2015).

45 Girod, C., Vitalis, R., Leblois, R. & Freville, H. Inferring population decline and expansion from microsatellite data: a simulation-based evaluation of the Msvar method. Genetics 188, 165–179, (2011).

46 Roman, J. & Palumbi, S. R. Whales before whaling in the North Atlantic. Science 301, 508–510, (2003).

47 Riccioni, G. et al. Spatio-temporal population structuring and genetic diversity retention in depleted Atlantic Bluefin tuna of the Mediterranean Sea. P Natl Acad Sci USA 107, 2102–2107, (2010).

48 Marra, A., Mona, S., Sa, R. M., D'Onghia, G. & Maiorano, P. Population Genetic History of Aristeus antennatus (Crustacea: Decapoda) in the Western and Central Mediterranean Sea. PLoS One 10, e0117272, (2015).

49 Pinsky, M. L. & Palumbi, S. R. Meta-analysis reveals lower genetic diversity in overfished populations. Mol Ecol 23, 29–39, (2014).

50 Li, H. & Durbin, R. Fast and accurate short read alignment with Burrows-Wheeler transform. Bioinformatics 25, 1754–1760, (2009).

51 Birol, I. et al. De novo transcriptome assembly with ABySS. Bioinformatics 25, 2872–2877, (2009).

52 McKenna, A. et al. The Genome Analysis Toolkit: a MapReduce framework for analyzing next-generation DNA sequencing data. Genome Res 20, 1297–1303, (2010).

53 Li, H. A statistical framework for SNP calling, mutation discovery, association mapping and population genetical parameter estimation from sequencing data. Bioinformatics 27, 2987–2993, (2011).

54 R: A Language and Environment for Statistical Computing (R Foundation for Statistical Computing, 2014).

55 Nei, M. Analysis of gene diversity in subdivided populations. Proc Natl Acad Sci U S A 70, 3321–3323, (1973).

56 Excoffier, L. & Lischer, H. E. Arlequin suite ver 3.5: a new series of programs to perform population genetics analyses under Linux and Windows. Mol Ecol Resour 10, 564–567, (2010).

57 Excoffier, L., Dupanloup, I., Huerta-Sanchez, E., Sousa, V. C. & Foll, M. Robust demographic inference from genomic and SNP data. PLoS Genet 9, e1003905, (2013).

58 Beaumont, M. A. in Simulation, Genetics, and Human Prehistory 135–154 (McDonald Institute for Archaeological Research, 2008).

59 Csillery, K., Francois, O. & Blum, M. G. B. abc: an R package for approximate Bayesian computation (ABC). Methods Ecol Evol 3, 475–479, (2012).

60 Gelman, A., Carlin, J., Stern, H. & Rubin, D. Bayesian Data Analysis. (CRC Press, 2004).

